# A Doppler Ultrasound Method to Measure Vascular Responses to Walking

**DOI:** 10.64898/2025.12.17.694736

**Authors:** Jose G Anguiano-Hernandez, Jenna K Burnett, Michael F Allen, Cody P Anderson, Kota Z Takahashi, Song-Young Park

## Abstract

Measuring leg blood flow (LBF) in response to walking can offer unique insights regarding mechanical and physiological functions of the leg. However, current gold-standard methods of measuring LBF are invasive. Alternatively, Doppler ultrasound is a promising method to noninvasively measure LBF. The purpose of this study was to demonstrate the utility of a novel ultrasound scanning technique to measure vascular reactivity to walking. Participants walked on a 0° slope, followed by a -5° and 5° slope in random order. A Doppler ultrasound probe was fastened over the dominant-side common femoral artery using a custom 3D printed bracket. LBF was recorded immediately before and after each walking condition. Baseline LBF was not significantly different before each walking condition (*p* = .577), but there were significant differences in post-walking (*p* < .001), with uphill causing the greatest increase in LBF. These results suggest that our ultrasound technique is repeatable during quiet standing and can detect the differences in blood flow across different walking intensities. Our method may be valuable to understand how the circulatory system responds to various locomotion tasks, and how cardiovascular disease may impair locomotor functions.

## INTRODUCTION

One of the main roles of the circulatory system is to transport oxygen and nutrients to metabolically active tissues. A large portion of tissues that are metabolically active during activities of daily living, like walking, are found in the leg muscles. The vascular system responds to the muscle’s metabolic demands during exercise by increasing oxygen delivery through a process called skeletal muscle hyperemia, or an increase in blood flow to active skeletal muscle. This increase in blood flow is proportional to the energy expended by the body [1–11]. As such, quantifying blood flow to the legs during walking may offer unique insights into how our bodies respond to various mechanical and metabolic demands. Assessing blood flow may also be valuable for understanding the effects of vascular disease on walking impairments, such as people with peripheral artery disease [12–23]. Therefore, measuring blood flow to the legs could elucidate the fundamental role of the circulatory system during locomotion, which may have broad impacts in basic and clinical research.

Estimating a leg’s blood flow (LBF) during locomotion is not trivial. Gold-standard methods of measuring peripheral blood flow include either the use of indwelling catheters inserted into the artery that supplies oxygen-rich blood to the leg in humans [4–10], or injecting biofluorescent microspheres into the circulation to track blood flow to individual muscles in animal models [1–3]. While these methods have greatly enhanced our understanding of the circulatory system’s response to exercise, they are highly invasive and should only be done by trained research or medical staff. In contrast, Doppler ultrasound offers an accurate [24,25] and noninvasive means of measuring LBF and the circulatory system’s response to exercise. However, such a technique has only been utilized to measure vascular responses to stationary exercises like passive leg movement [26,27], isolated knee extensions [28], and cycling [29,30]. Measuring vascular responses during walking with Doppler ultrasound is challenging due to repeated hip flexion and extension combined with the body’s natural oscillations. One solution to this challenge could be to affix a Doppler ultrasound probe over a person’s common femoral artery, such that LBF measurements can be made immediately before and after a walking bout. Such a technique could give insight into how the leg’s energy demands change with different walking intensities.

The purpose of this study was to establish a novel ultrasound technique by comparing the leg hyperemia changes induced by sloped walking as a testing framework. Relative to walking on a level slope, walking on an incline or decline slope increases or reduces the metabolic cost of walking, respectively [31]. The metabolic demand of an exercise task is met by increased blood flow to metabolically active tissue [4–10], allowing us to test our ultrasound method’s sensitivity to detect differences in leg hyperemia across a range of walking demands. First, we hypothesized that our ultrasound scanning technique will demonstrate good repeatability by finding similar LBF before walking conditions. Second, we hypothesized that our ultrasound technique will be sensitive enough to detect significant differences in the change in LBF between walking conditions. As an exploratory analysis, we quantified LBF Recovery Time (LBF-RT), or the time required for LBF to return to its resting state after walking.

## MATERIALS AND METHODS

### Participants

Pilot testing of our experiment found an effect size f = .655 when comparing blood flow velocity during walking on a -5°, 0°, and 5° slope in 4 healthy young adults. A priori power analysis (G*Power 3.1) suggested a sample size of 6 participants. To ensure sufficient statistical power, 14 healthy young adults (8 females, 27.1 ± 4.7 years old, 67.4 ± 8.6 kg, 1.7 ± 0.09 m) were recruited. Informed consent was obtained from all participants before data collection in accordance with the University of Utah Institutional Review Board (IRB #155345). Exclusion criteria included having a leg injury or fracture within the last 6 months, a neurological disorder of the legs (e.g., peripheral neuropathy, stroke), a vascular disease (e.g., peripheral artery disease, diabetes), taking medication that causes dizziness, or the inability to walk without an assistive device (i.e., cane or walker). Participants were asked to avoid rigorous exercise, alcohol, caffeine, large meals, and tobacco products at least 4 hours prior to the data collection.

### 3D Printed Bracket Design

To design the 3D printed bracket, the ultrasound probe was 3D scanned (HandySCAN 3D, Creaform, Canada) and the model was imported into TinkerCAD (Autodesk, San Francisco, CA, USA). An ovular prism was superimposed over the probe model and the shape of the probe was subtracted. Four rectangular prisms were placed at an angle relative to the probe’s orientation to facilitate fastening the bracket to the participant. The model was split in half and attachment fixtures were added to allow donning and doffing of the bracket over the probe. The bracket was printed in PLA filament with a Prusa i3 Mk3S+ 3D printer (Prusa Research, Prague, Czech Republic) at 30% infill (Figure 1A). The 3D print was sanded at the attachment fixtures to enable donning and doffing of the bracket over the probe (Figure 1B).

**Figure 1.**
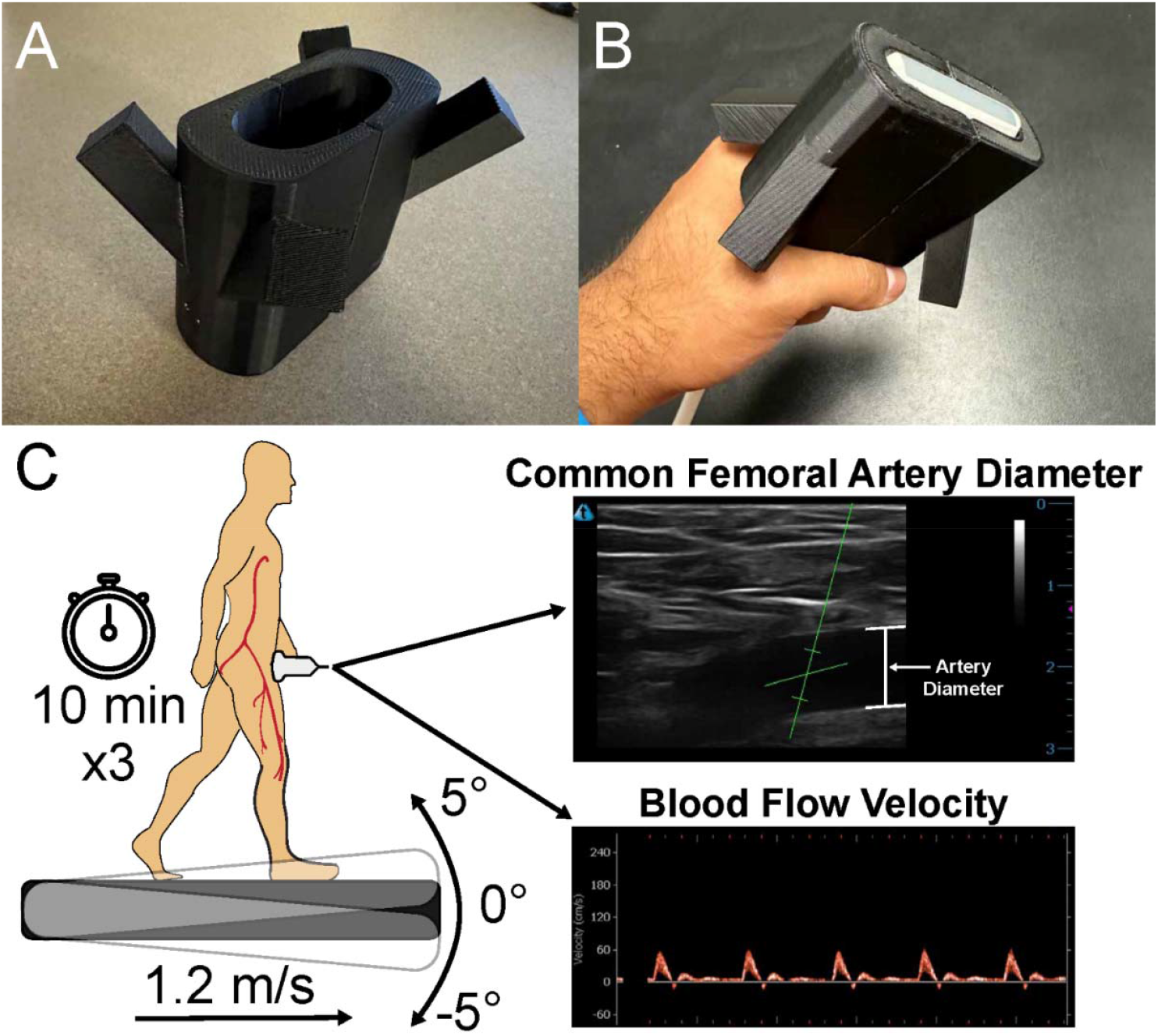
A) The 3D printed bracket used to fasten the ultrasound probe over the participant’s femoral artery, B) the bracket placed on the ultrasound probe, C) illustration of our experimental protocol. Participants walked on a 0° slope first, then a -5° and 5° slope in random order for 10 minutes each at 1.2 m/s.

### Experimental Protocol

Data collection took place at the Advanced Rehabilitation Technology Studio in the Craig H. Neilsen Rehabilitation Hospital at the University of Utah. Upon arrival, participants donned custom compression shorts that allowed the investigators to place a Doppler ultrasound probe (uSmart 3300 NexGen and 15L4A Linear Transducer, Terason, Burlington, MA, USA) over the common femoral artery (inguinal area) of the participant’s dominant leg (Figure 1C). We chose to scan the common femoral artery because it is the largest artery that carries oxygen-rich blood to the legs. The ultrasound probe was fixed to the participant using a custom 3D printed bracket and athletic tape. Imaging depth of the B-Mode image was adjusted such that the femoral artery remained central in the frame, typically either 3 or 4 cm. The insonation angle of the ultrasound beam was set to 60° [32]. The advantage of fixing the ultrasound probe over the participant’s common femoral artery while they walk is the time saved collecting post-walking leg blood flow measurements. If we did not fix the probe to the participant, it would require some time to find an adequate image of the common femoral artery after walking due to maneuvering the participant’s clothes to allow for probe placement, and the actual scanning for the femoral artery. If too much time has elapsed after the end of a walking bout, much of the vascular response to the walking bout will be lost, causing inaccurate leg blood flow measurements. Our method of fastening the probe to the participant bypasses the need for moving the participant’s clothes and searching for the artery, which allowed us to begin recording the vascular response to walking just seconds after a walking bout.

After placing and fixing the ultrasound, participants rested seated with their legs extended and supported for 15 minutes before each walking condition. Participants walked on a treadmill (FIT5, Bertec, Columbus, OH, USA) at 1.2 m/s on a 0° slope for 10 minutes in the first condition, followed by a 5° and -5° slope in random order at the same walking speed and duration while ultrasound videos were recorded. The 0° slope condition was completed first so participants could acclimate to walking with the ultrasound probe fixed over their inguinal region. A 30-second baseline measurement of femoral artery diameter and blood flow velocity were collected before each walking condition. Participants stepped off the treadmill immediately after each walking condition and a 5-minute recovery ultrasound measurement was collected. Participants then rested for at least 15 minutes before the next walking condition began.

### Data Analysis

Ultrasound videos were trimmed into 30-second baseline and 5-minute post-walking videos. FloWaveUS, a validated open-source software, extracted instantaneous femoral artery diameter and blood flow velocity time series from ultrasound videos as a time series [33]. The software digitizes the B-Mode image and Doppler velocity graph from duplex ultrasound videos and converts pixel distances to cm and cm/s, respectively. Artery diameter data were filtered using a 2^nd^ order recursive low-pass Butterworth filter with a 10 Hz cutoff frequency, as determined by residual analysis [34]. A range of cutoff frequencies were tested, and the best cutoff frequency was selected based on the optimization of the residuals between the raw and filtered data [34] (Supplementary Figure S1A). We chose a low-pass filter rather than a high-pass filter to preserve the low frequency changes in artery diameter that occur during the cardiac cycle, while eliminating high frequency changes in diameter that are not physiological. Blood flow velocity data were filtered using a 5^th^ order median filter (Supplementary Figure S1B). We used a moving median filter for the raw blood flow velocity data because it is better suited to remove noise that presents randomly throughout a signal. Filtered artery diameter and blood flow velocity time series data were used to calculate LBF in L/min using Equation 1. A 30-second average of pre-walking artery diameter, blood flow velocity, and LBF were used to compare baseline values between conditions. Pre-walking values were subtracted from post-walking values to calculate and compare the change in each variable across walking conditions.

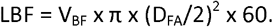

Equation 1: Leg blood flow (LBF) calculated as a function of blood flow velocity (V_BF_) and femoral artery diameter (D_FA_). V_BF_ is expressed in cm/s and D_FA_ is expressed in cm.

We quantified LBF-RT by calculating a condition-specific threshold that was equal to the 30-second baseline LBF average plus one half of a standard deviation. The post-walking LBF time series was then time-averaged into 2.5 second averages (Figure 2). To smooth the time series and prevent multiple crossings of the threshold, a 3-sample moving mean of the time-averaged LBF time series data was calculated. The point when the time-average recovery time series crossed the threshold was identified and labeled as the LBF-RT. All filtering and calculations were performed using custom MATLAB scripts (2023a, The Mathworks, Natick, MA, USA).

**Figure 2.**
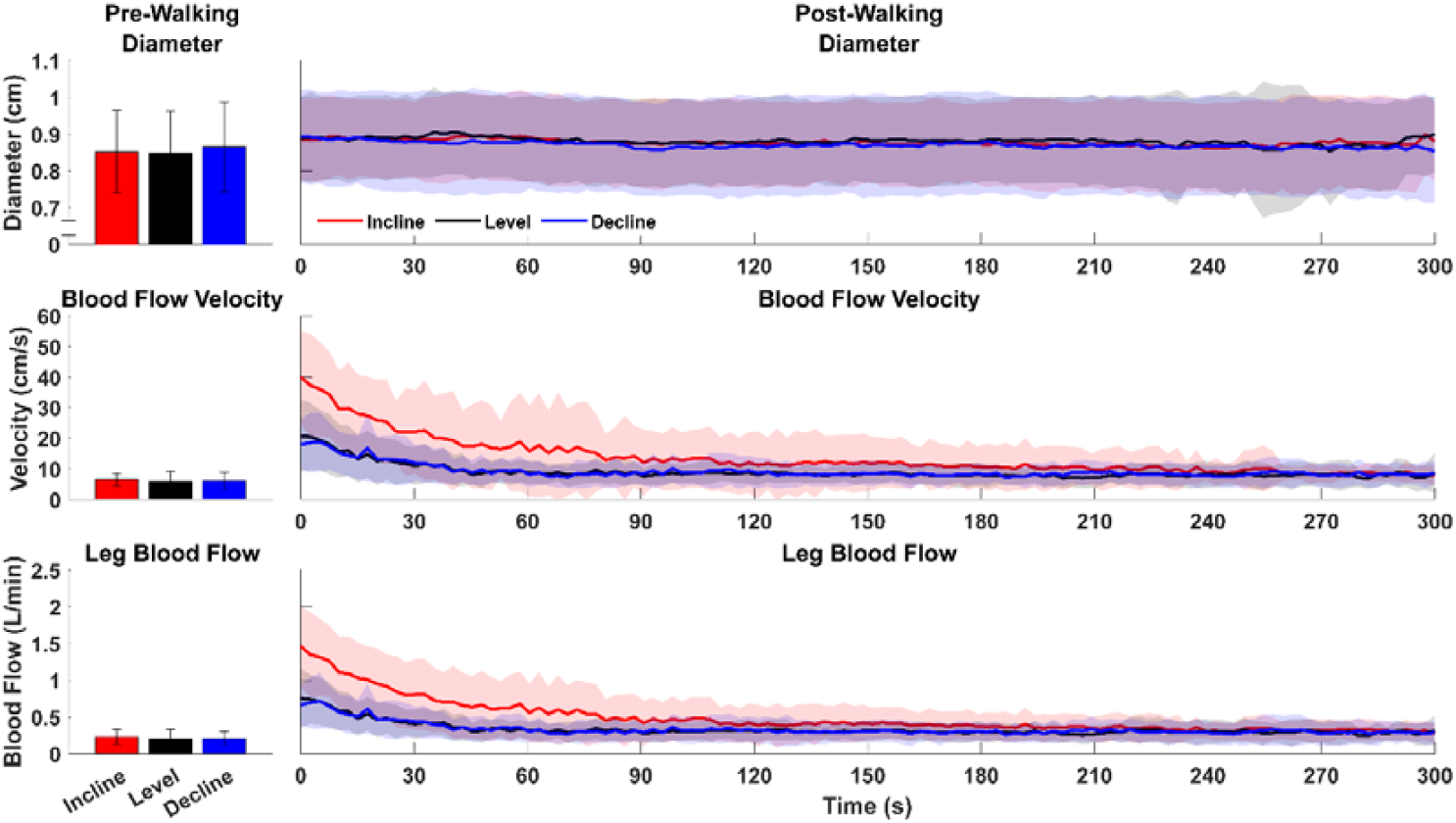
Baseline (left column) and post-walking (right column) Femoral Artery Diameter, Blood Flow Velocity, And Leg Blood Flow across walking conditions. Baseline values are 30-second averages during the baseline ultrasound measurement. Recovery time series data points are 2.5 second averages of the filtered data. Shaded regions around the mean time series are one standard deviation.

### Statistical Analysis

SPSS 30 (IBM, Armonk, NY, USA) was used for statistical analyses. Shapiro-Wilks tests were used to check data for normality. To assess the repeatability of our technique, separate one-way repeated measures ANOVA were used to compare baseline artery diameter, blood flow velocity, and LBF across slope conditions. To assess the sensitivity of our technique, we used separate one-way repeated measures ANOVA to compare the change in artery diameter, blood flow velocity, and LBF from baseline across conditions. For our exploratory analysis, a repeated measures ANOVA was used to compare LBF-RT across walking conditions. Main effects were investigated using Bonferroni-corrected pairwise comparisons. All statistical tests were performed using an α = 0.05.

## RESULTS

### Femoral Artery Diameter

Baseline artery diameter was similar across walking conditions (Incline: .85 ± .11 cm, Level: .85 ± .11 cm, Decline: .87 ± .12 cm, p = .297), suggesting that our ultrasound method can be used to repeatedly measure femoral artery diameter. The change in artery diameter from baseline was similar across walking conditions (Incline: .037 ± .052 cm, Level: .044 ± .050 cm, Decline: .020 ± .39 cm, *p* = .316).

### Blood Flow Velocity

Baseline blood flow velocity was similar across walking conditions (Incline: 6.36 ± 2.11 cm/s, Level: 5.91 ± 3.31 cm/s, Decline: 6.029 ± 2.72 cm/s, *p* = .680), suggesting that our ultrasound method can repeatedly measure femoral artery blood flow velocity. The change in blood flow velocity was different across slopes (p < .001), with uphill walking increasing blood flow velocity more than the level walking (23.38 ± 12.62 vs 9.29 ± 8.29 cm/s, p_adj_ < .001) and downhill walking (23.38 ± 12.62 vs 9.36 ± 5.89 cm/s, p_adj_ < .001).

### Leg Blood Flow

Baseline LBF was similar across walking conditions (Incline: .23 ± .099 L/min, Level: .20 ± .13 L/min, Decline: .21 ± .094 L/min, p = .525), suggesting that our ultrasound method can repeatedly measure LBF. The change in LBF was different across slopes (*p* < .001), with uphill walking increasing blood flow velocity more than the level walking (.87 ± .44 vs .35 ± .28 L/min, *p*_*adj*_ = .001) and downhill walking (.87 ± .44 vs .35 ± .19 L/min, *p*_*adj*_ < .001).

### Leg Blood Flow Recovery Time

One participant’s LBF did not return to baseline value during the post-walking measurement. This participant had very low LBF before one walking condition, resulting in a very low LBF-RT threshold. This participant did not reach that threshold after the walking condition, so this participant was excluded from the ANOVA. LBF-RT was different across walking slopes (*p* = .007). LBF-RT was greater after walking uphill compared to level (94.42 ± 78.37 vs 39.04 ± 29.057 s, *p*_*adj*_ = .021) and between the incline and decline conditions (94.42 ± 78.37 s vs 30.39 ± 16.36 s, *p*_*adj*_ = .027), suggesting that it takes longer for LBF to return to baseline after walking uphill compared to walking on a level and downhill surface (Figure 4).

## DISCUSSION

We sought to establish an ultrasound technique to noninvasively measure vascular reactivity to walking. We found similar baseline artery diameter, blood flow velocity, and LBF values before each walking condition, supporting our first hypothesis that our method is repeatable. Our second hypothesis was partially supported, as only changes in blood flow velocity and LBF were significantly different across walking conditions. Our findings demonstrate the utility of our method for non-invasively measuring vascular reactivity to walking. These results also suggest that our ultrasound method could be utilized to investigate the circulatory system’s role in human locomotion.

Our ultrasound technique showed good repeatability during pre-walking LBF measurements. We found no significant differences between conditions in our outcome variables before walking, suggesting that our ultrasound method is repeatable during quiet standing. This result is likely due to fastening the ultrasound probe over the participant’s common femoral artery during placement, which ensures that we are recording ultrasound images of the same section of the artery before each walking condition. Based on the similar values of our outcome variables at baseline, we also speculate that the ultrasound probe is not displacing significantly during walking bouts, walking periods, or any transitions between walking and resting.

Our ultrasound method was also sensitive enough to detect significant differences in blood flow velocity and LBF induced by various walking slopes, which are slopes that are known to vary mechanical and physiological walking demands [31,35–42]. Our results suggest that leg hyperemia could be more related to the increase in blood flow velocity induced by walking instead of the change in femoral artery diameter. Recall that our LBF calculation is a function of artery diameter and blood flow velocity (Equation 1). Because there were significant differences in the increase in blood flow velocity between conditions, and no significant differences in vasodilation between conditions (Figure 3), we can conclude that the increase in leg blood flow was mainly driven by increased blood flow velocity. In future studies, our technique could aid in elucidating the energetic and cardiovascular underpinnings of walking function in those with and without cardiovascular disease by allowing investigators to non-invasively investigate the circulatory system’s role in locomotion.

**Figure 3.**
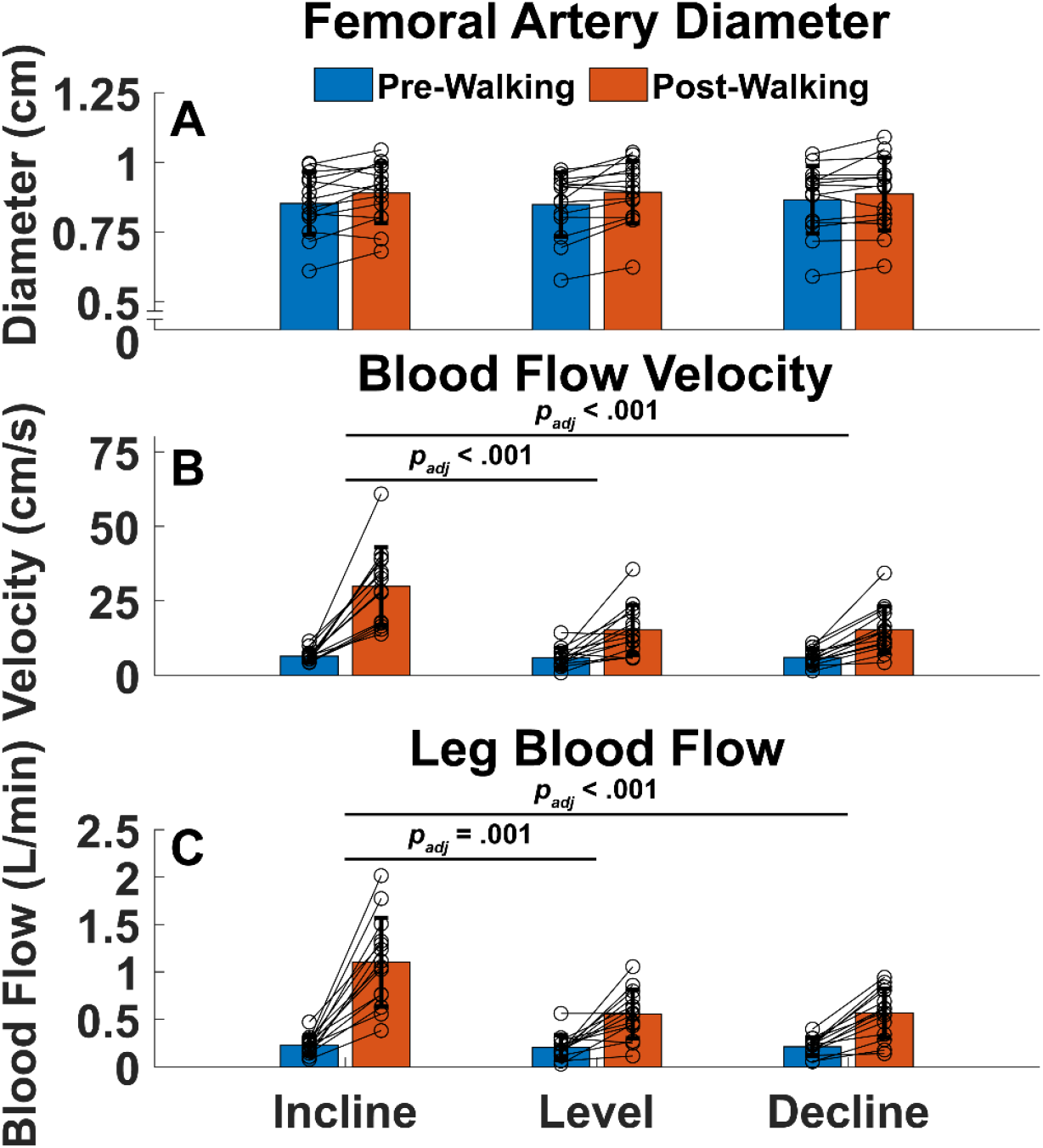
Thirty second pre- and post-walking A) Femoral Artery Diameter, B) Blood Flow Velocity, and C) Leg Blood Flow. Connected circles are individual participant data. Femoral artery dilation was not significantly different across walking conditions. Incline walking caused the greatest change from rest in blood flow velocity and leg blood flow. Bars are mean ± 1 standard deviation. Brackets connecting groups of bars denote significantly different changes in variable being plotted (post minus pre-walking) between walking conditions.

As an exploratory aim, we quantified LBF-RT after each walking condition. Our results indicate LBF takes 30 to 94 seconds to return to baseline in healthy young adults, depending on walking slope (Figure 4). Because the probe was fastened to the participant and stayed in place throughout the data collection, we were able to immediately record LBF when the participant stepped off the treadmill. However, if the probe had to be replaced to find the common femoral artery post-walking, a significant portion of the LBF response may be missed and result in LBF measurements that do not truly reflect the vascular system’s response to walking, particularly since LBF begins to decrease immediately after walking (Figure 2). These results emphasize the importance of recording post-walking LBF as soon as possible after walking. Our method of affixing the ultrasound probe to the participant ensures that we can measure leg hyperemia immediately after a walking bout to record the most accurate vascular response possible.

**Figure 4.**
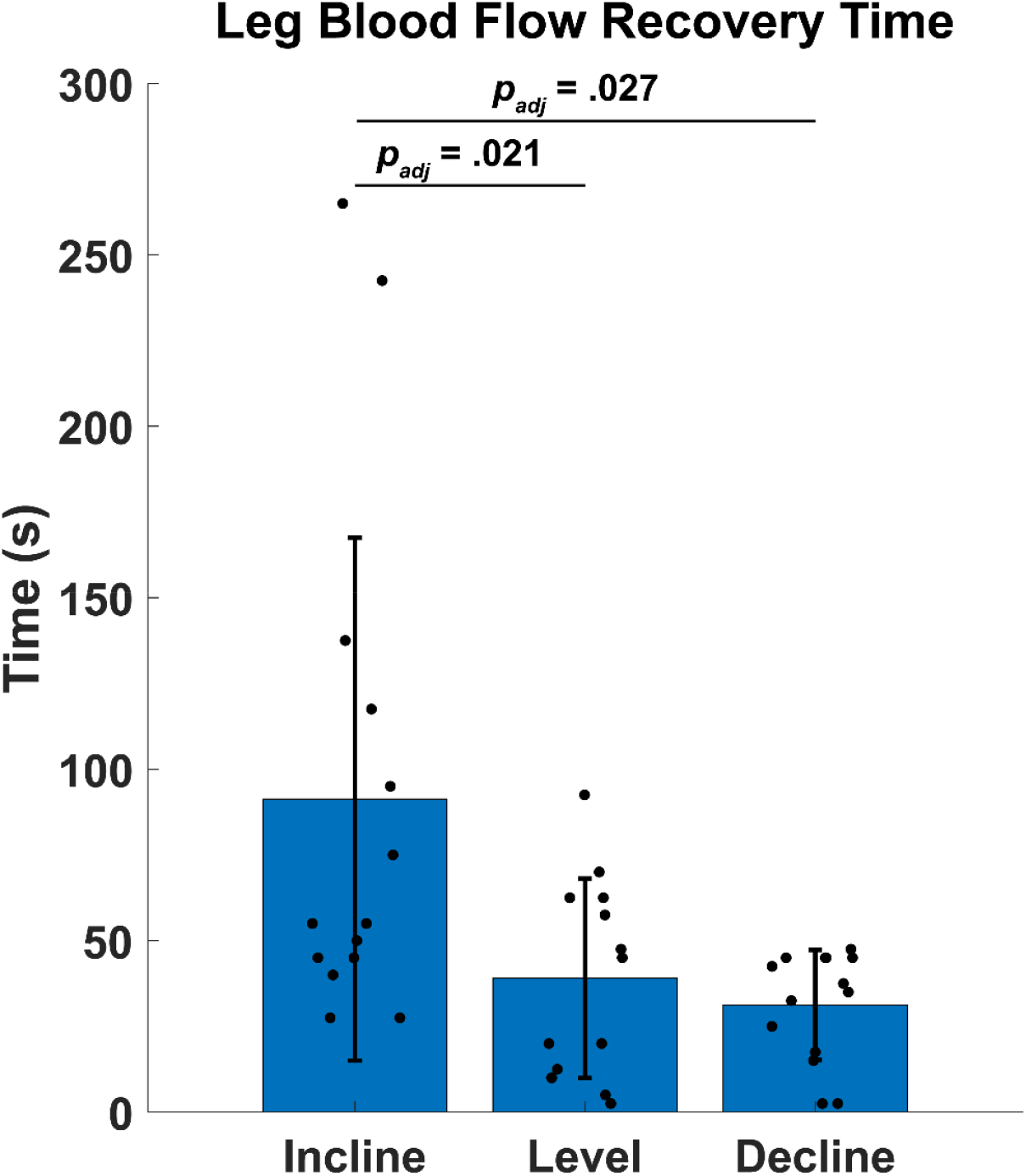
Leg Blood Flow Recovery Time for all walking conditions. Incline walking resulted in the greatest Leg Blood Flow Recovery Time. Horizontal lines denote significant pairwise differences in Leg Blood Flow Recovery Time. Bars are mean ± 1 standard deviation. Filled circles are individual participant data (n=13). Brackets connecting bars represent significantly different pairwise comparisons.

There are several limitations of the current study, including the absence of ultrasound images during walking, metabolic or central hemodynamic measures, and comparison with gold-standard methods. Analyzing LBF non-invasively during walking could paint a more complete picture of vascular reactivity to walking, but repeated hip flexion and extension prevented analysis of femoral artery ultrasound videos. To circumvent this limitation, we fastened the ultrasound probe over the participant’s common femoral artery so we can record LBF immediately after walking. While metabolic and central hemodynamic measures like indirect calorimetry or heart rate were not included in this study, we speculate that our non-invasive LBF estimates would show positive correlations with those measures. For instance, our study found that increase in LBF was about 2.5 times greater (.87 vs .35 L/min) after walking on a 5° slope compared to a level slope (Figure 3C), which closely aligns with a previous study reporting that the metabolic cost of transport during walking on a 5.7° slope was 2.75 times greater when compared to walking on a level slope [31]. Furthermore, we speculate that our non-invasive LBF estimates may agree well with those derived from more invasive methods. For example, in a study using indwelling catheters[5], they reported a peak LBF of 4.15 L/min during walking on a 2.86° incline at 1.39 m/s. In our experiment, we found an average peak LBF of 3.96 L/min after walking on a 5° incline at 1.2 m/s (Supplementary Figure S2). Similar peak LBF results between the two studies, albeit at slightly different walking intensities, suggest that our non-invasive estimates may yield similar results to more invasive methods. Our present study motivates future studies to compare our non-invasive LBF measures with previously validated techniques or metabolic or central hemodynamics measures. Such efforts will enable greater understanding of the relationship between metabolic cost of walking and skeletal muscle blood flow.

Other limitations of this study include not assessing the intra- or inter-operator variability of our ultrasound method. All the LBF measurements in this study were done by the same investigator, so showing that our results could be replicated by multiple investigators would demonstrate our technique’s utility in research. Additionally, demonstrating the repeatability of our method within participants on different days may add to our method’s utility in future research. However, these limitations did not influence the outcome of the present study because of its single-visit, within-subjects design.

## DATA AVAILABILITY

The datasets used and/or analysed during the current study available from the corresponding author on reasonable request.

## ACKNOWLEDGMENTS

We would like to thank Shawn Plucinski and Charles Fisher in the University of Nebraska at Omaha Criss Library Creative Production Lab for their assistance with 3D printing. We also thank Kylee Heap, Chelsea Haddy, and Emma Caringella for their assistance with data collection.

## AUTHOR CONTRIBUTIONS

All authors conceived and designed research; J.G.A and J.K.B performed experiments, J.G.A analyzed data, all authors interpreted results of experiments, J.G.A prepared figures, J.G.A drafted manuscript, all authors edited and revised manuscript, all authors approved final version of manuscript.

## SUPPLEMENTAL MATERIAL

**Supplementary Figure S1:**
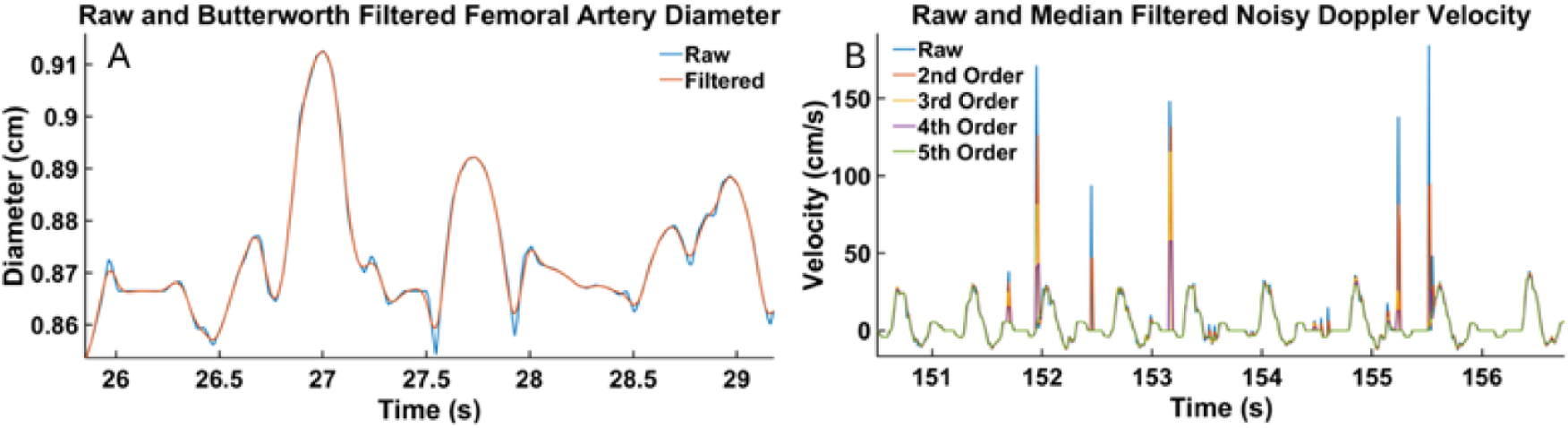
Representative plots of raw and filtered A) femoral artery diameter and B) blood flow velocity time-series data from one walking condition. Femoral artery diameter was filtered with a 2^nd^ order recursive Butterworth filter with a 10 Hz cutoff frequency. Blood flow velocity data was filtered using a 5^th^ order 1-dimensional median filter. A 5^th^ order median filter (green line) was best for reducing noise without affecting the signal.

**Supplementary Figure S2:**
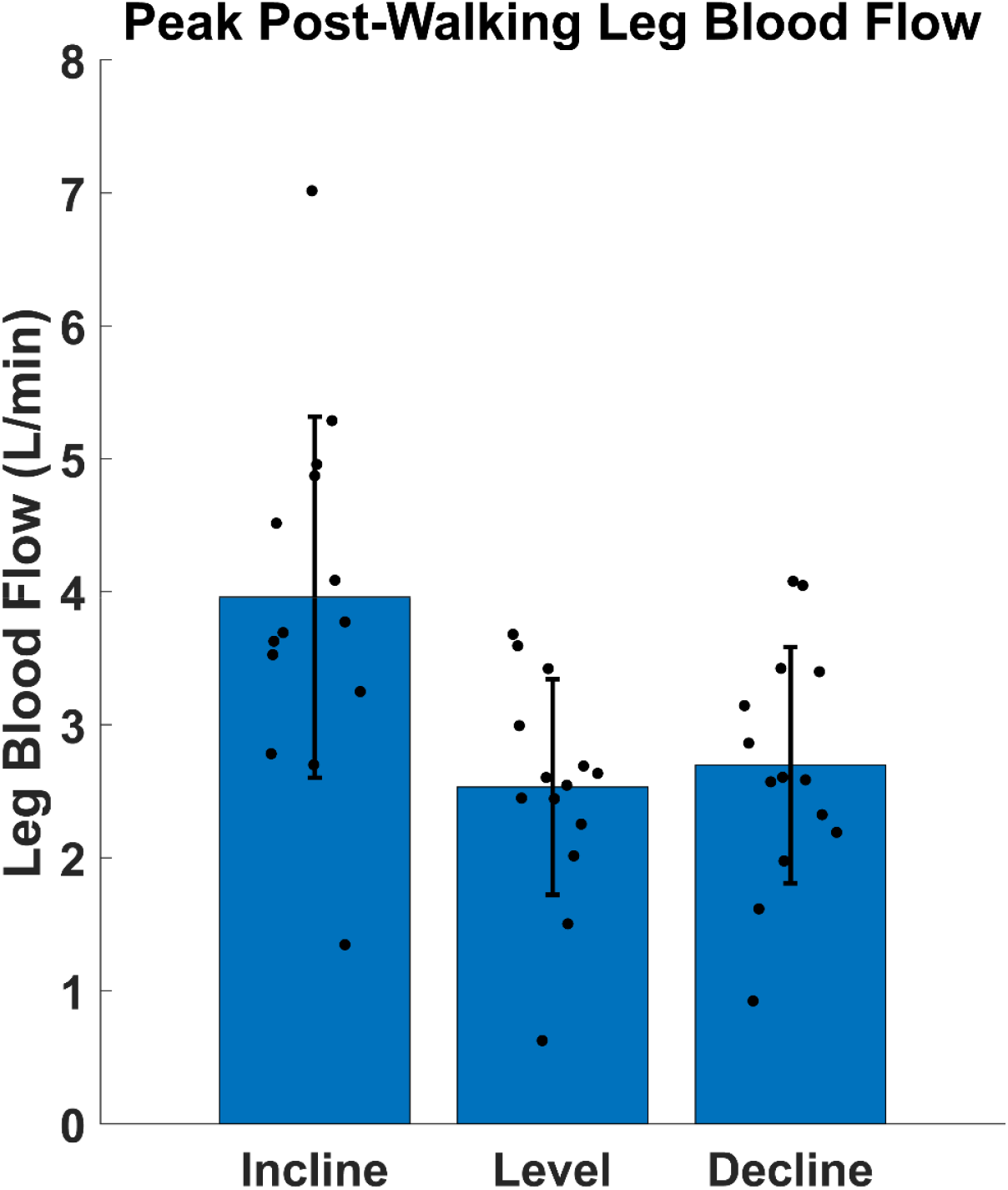
Peak Leg Blood Flow in the first 20 seconds after each walking condition. Incline walking resulted in the greatest peak Leg Blood Flow compared to Level and Decline walking.

## FUNDING

National Institutes of Health, National Institute of Child Health and Human Development, Project Number: 1R01HD106911-01A1 (to KZT and SP).

National Institutes of Health, National Institute of Diabetes and Digestive and Kidney Diseases, Project Number: 1T32DK11096601 (to JKB).

## ETHICS STATEMENT

All procedures involving human participants were approved by the University of Utah Institutional Review Board and conducted in accordance with the ethical standards of the Declaration of Helsinki. Written informed consent was obtained from all participants prior to their inclusion in the study.

## REFERENCES

1. Rubenson, J., Henry, H. T., Dimoulas, P. M. & Marsh, R. L. The cost of running uphill: linking organismal and muscle energy use in guinea fowl (Numida meleagris). Journal of Experimental Biology 209, 2395–2408 (2006).

2. Marsh, R. L. & Ellerby, D. J. Partitioning locomotor energy use among and within muscles Muscle blood flow as a measure of muscle oxygen consumption. Journal of Experimental Biology 209, 2385–2394 (2006).

3. Marsh, R. L., Ellerby, D. J., Carr, J. A., Henry, H. T. & Buchanan, C. I. Partitioning the Energetics of Walking and Running: Swinging the Limbs Is Expensive. Science 303, 80–83 (2004).

4. Calbet, J. A. L. et al. Cardiac output and leg and arm blood flow during incremental exercise to exhaustion on the cycle ergometer. Journal of Applied Physiology 103, 969–978 (2007).

5. Leahy, M. G. et al. Assessing Leg Blood Flow and Cardiac Output During Running Using Thermodilution. Scandinavian Journal of Medicine & Science in Sports 34, e14705 (2024).

6. Andersen, P. & Saltin, B. Maximal perfusion of skeletal muscle in man. The Journal of Physiology 366, 233–249 (1985).

7. Proctor, D. N. et al. Leg blood flow during submaximal cycle ergometry is not reduced in healthy older normally active men. Journal of Applied Physiology 94, 1859–1869 (2003).

8. Proctor, D. N., Koch, D. W., Newcomer, S. C., Le, K. U. & Leuenberger, U. A. Impaired leg vasodilation during dynamic exercise in healthy older women. Journal of Applied Physiology 95, 1963–1970 (2003).

9. Proctor, D. N. et al. Leg Blood Flow and VO2 during Peak Cycle Exercise in Younger and Older Women. Medicine & Science in Sports & Exercise 36, 623 (2004).

10. Proctor, D. N. et al. Reduced leg blood flow during dynamic exercise in older endurance-trained men. Journal of Applied Physiology 85, 68–75 (1998).

11. Laughlin, M. H. Cardiovascular response to exercise. Am J Physiol 277, S244–259 (1999).

12. Koutakis, P. et al. Abnormal Joint Powers Before And After The Onset Of Claudication Symptoms. J Vasc Surg 52, 340–347 (2010).

13. McDermott, M. M. et al. Asymptomatic Peripheral Arterial Disease Is Associated With More Adverse Lower Extremity Characteristics Than Intermittent Claudication. Circulation 117, 2484–2491 (2008).

14. Chen, S.-J. et al. Bilateral claudication results in alterations in the gait biomechanics at the hip and ankle joints. Journal of Biomechanics 41, 2506–2514 (2008).

15. McDermott, M. M. EXERCISE TRAINING FOR INTERMITTENT CLAUDICATION. J Vasc Surg 66, 1612–1620 (2017).

16. McDermott, M. M. Functional Impairment in Peripheral Artery Disease and How to Improve It in 2013. Curr Cardiol Rep 15, 347 (2013).

17. Szymczak, M., Krupa, P., Oszkinis, G. & Majchrzycki, M. Gait pattern in patients with peripheral artery disease. BMC Geriatr 18, 52 (2018).

18. Koutakis, P. et al. Joint torques and powers are reduced during ambulation for both limbs in patients with unilateral claudication. J Vasc Surg 51, 80–88 (2010).

19. Wurdeman, S. R. et al. Patients with peripheral arterial disease exhibit reduced joint powers compared to velocity-matched controls. Gait Posture 36, 506–509 (2012).

20. Rahman, H. et al. Peripheral Artery Disease Causes Consistent Gait Irregularities Regardless of the Location of Leg Claudication Pain. Ann Phys Rehabil Med 67, 101793 (2024).

21. McDermott, M. M. et al. PROGNOSTIC VALUE OF FUNCTIONAL PERFORMANCE FOR MORTALITY IN PATIENTS WITH PERIPHERAL ARTERIAL DISEASE. J Am Coll Cardiol 51, 1482–1489 (2008).

22. Grenon, S. M. et al. Walking Disability in Patients with Peripheral Artery Disease is Associated with Arterial Endothelial Function. J Vasc Surg 59, 1025–1034 (2014).

23. White, S. H. et al. Walking performance is positively correlated to calf muscle fiber size in peripheral artery disease subjects, but fibers show aberrant mitophagy: an observational study. Journal of Translational Medicine 14, 284 (2016).

24. Hoyt, K., Hester, F. A., Bell, R. L., Lockhart, M. E. & Robbin, M. L. Accuracy of Volumetric Flow Rate Measurements. J Ultrasound Med 28, 1511–1518 (2009).

25. Zierler, B. K. et al. Accuracy of duplex scanning for measurement of arterial volume flow. Journal of Vascular Surgery 16, 520–526 (1992).

26. Trinity, J. D. et al. Limb movement-induced hyperemia has a central hemodynamic component: evidence from a neural blockade study. Am J Physiol Heart Circ Physiol 299, H1693–H1700 (2010).

27. Park, S.-Y. et al. Effects of passive and active leg movements to interrupt sitting in mild hypercapnia on cardiovascular function in healthy adults. J Appl Physiol (1985) 132, 874–887 (2022).

28. Rossman, M. J. et al. Oral antioxidants improve leg blood flow during exercise in patients with chronic obstructive pulmonary disease. Am J Physiol Heart Circ Physiol 309, H977–H985 (2015).

29. Ratchford, S. M. et al. Salt restriction lowers blood pressure at rest and during exercise without altering peripheral hemodynamics in hypertensive individuals. Am J Physiol Heart Circ Physiol 317, H1194–H1202 (2019).

30. Cohen, J. N., Sole, R. T., Zafiris, E. & Au, J. S. Efficacy of a hands-free vascular ultrasound probe holder in active and inactive limbs during cycling exercise. Journal of Applied Physiology 138, 389–396 (2025).

31. Minetti, A. E., Moia, C., Roi, G. S., Susta, D. & Ferretti, G. Energy cost of walking and running at extreme uphill and downhill slopes. Journal of Applied Physiology 93, 1039–1046 (2002).

32. Logason, K. et al. The Importance of Doppler Angle of Insonation on Differentiation Between 50–69% and 70–99% Carotid Artery Stenosis. European Journal of Vascular and Endovascular Surgery 21, 311–313 (2001).

33. Coolbaugh, C. L., Bush, E. C., Caskey, C. F., Damon, B. M. & Towse, T. F. FloWave.US: validated, open-source, and flexible software for ultrasound blood flow analysis. Journal of Applied Physiology 121, 849–857 (2016).

34. Winter, D. Biomechanics and Motor Control of Human Movement. (John Wiley & Sons, Ltd, 2009). doi:10.1002/9780470549148

35. Lay, A. N., Hass, C. J., Richard Nichols, T. & Gregor, R. J. The effects of sloped surfaces on locomotion: An electromyographic analysis. Journal of Biomechanics 40, 1276–1285 (2007).

36. Franz, J. R. & Kram, R. The effects of grade and speed on leg muscle activations during walking. Gait & Posture 35, 143–147 (2012).

37. Franz, J. R. & Kram, R. How does age affect leg muscle activity/coactivity during uphill and downhill walking? Gait Posture 37, 378–384 (2013).

38. Abe, D., Fukuoka, Y. & Horiuchi, M. Economical Speed and Energetically Optimal Transition Speed Evaluated by Gross and Net Oxygen Cost of Transport at Different Gradients. PLoS One 10, e0138154 (2015).

39. Alexander, N. & Schwameder, H. Effect of sloped walking on lower limb muscle forces. Gait & Posture 47, 62–67 (2016).

40. Alexander, N., Strutzenberger, G., Ameshofer, L. M. & Schwameder, H. Lower limb joint work and joint work contribution during downhill and uphill walking at different inclinations. Journal of Biomechanics 61, 75–80 (2017).

41. Nuckols, R. W. et al. Mechanics of walking and running up and downhill: A joint-level perspective to guide design of lower-limb exoskeletons. PLoS One 15, e0231996 (2020).

42. Papachatzis, N. & Takahashi, K. Z. Mechanics of the human foot during walking on different slopes. PLOS ONE 18, e0286521 (2023).

